# Competition and Integration of Visual and Goal Vector Signals for Spatial Navigation

**DOI:** 10.1101/2024.05.14.594206

**Authors:** Sandhiya Vijayabaskaran, Sen Cheng

## Abstract

Integrating different sources of information is essential to successful spatial navigation. For instance, animals must often rely on a combination of visual impressions, self-motion, olfaction, and other signals to navigate to a goal. This is especially important when navigating in uncertain environments, where switching from one source of information to another or integrating multiple sources of information may be required to make navigation decisions. We propose a computational model of the interaction of visual and goal-vector signals based on reinforcement learning and use it to study behavior and spatial representations. Our model demonstrates that the ability to navigate using each information source independently, in addition to integrating them, is crucial to successfully navigating in uncertain environments. Counterintuitively, our model also shows that when one of the signals is removed, navigation may be improved if the remaining signal is reliable and sufficient to navigate, however, this improvement comes at the expense of robustness.

## Introduction

Animals must make use of several available sources of information and take into account various sensory modalities to navigate: visual, self-motion, odor, tactile, magnetosensory perception, and so on could all be used by animals to navigate their surroundings (Parra-Barrero et al., 2023). The redundancy of information encoded by these input signals enables them to navigate effectively in the face of uncertainty and in changing environments. However, successfully integrating different information sources to form a coherent representation of the environment is far from straightforward. The reliability and accuracy of each source can vary, and animals must constantly evaluate and prioritize which information to rely on in any given situation and also be able to deal with conflicting information.

In neuroscience and behavioral research, this issue has been primarily studied in the context of how vision and self-motion information interact to influence navigation and spatial representations. The self-motion signal is presumably used to compute and update a vector representation of the goal (or goal vector). Behavioral studies show that when animals are navigating using visual cues, they still manage to arrive at the goal when the lights are turned off at the start of, or during, navigation (Collett et al., 1986; Save, 1997). Similar behavior is observed in humans, where participants are able to navigate blindfolded to a previously seen goal (Mittelstaedt & Mittelstaedt, 2001), demonstrating that both vision and self-motion can be used to navigate to the goal.

While it is clear that these two signals interact, the precise nature of this interaction remains unknown. One question is whether the two signals cooperate or compete — are both signals somehow integrated to determine which direction to move, or is one signal selected over the other based on some criteria (Parra-Barrero et al., 2023)? In the case of the former, it is also unclear how and to what extent each of the signals contributes to the integration, and in the latter, which signal is chosen and how. Experimentally, both outcomes have been observed in behavioral studies in rodents: an integration of both signals, with visual cues dominating, as well as a selection of one of them (Etienne & Jeffery, 2004; Etienne et al., 1996; Alyan & Jander, 1994; Etienne et al., 1990). Both kinds of interactions have also been observed in behavioral experiments in humans. While some studies show that humans combine vision and self-motion to navigate (Xu et al., 2017; X. Chen et al., 2017), others suggest that one or the other is preferred (Foo et al., 2005; Zhao & Warren, 2015). In general, the behavioral evidence is consistent with the integration of signals when the mismatch between them is small and selection when the mismatch is intermediate or large.

Neuroscientists, on the other hand, have focused on linking place cell firing to these information sources. Again, both visual cues and self-motion have been shown to influence place cell firing, although to different extents. On one hand, multiple studies have shown that place cells consistently fire at a fixed distance from the start location of a linear track, even when the position of the start box is shifted (Gothard et al., 1996; Redish et al., 2000; Bjerknes et al., 2018). However, various visual cues, such as the position of a cue card, the introduction of barriers, and environmental borders, have also been shown to strongly modulate the location of place cell firing (Muller & Kubie, 1987; O’Keefe & Burgess, 1996).

The observed differences may be due to the difficulty of fully dissociating the two signals in experiments, but a few studies have explicitly addressed this question using lesions (Stackman et al., 2002), virtual reality (G. Chen et al., 2013), and genetically modified animals (Rochefort et al., 2011). Although these studies clearly demonstrate that both vision and self-motion jointly impact place cell firing, they reach different conclusions regarding the relative contributions of the two, which could potentially be attributed to the different types of interventions involved. This disparity in experimental findings is also reflected in computational models of place cells, which make different assumptions about the underlying mechanisms. While some models are based on the assumption that self-motion information drives place cell firing (Samsonovich & McNaughton, 1997; McNAUGHTON et al., 1996), others propose that visual cues play a predominant role (Franzius et al., 2007; Burgess et al., 2000; Sharp et al., 1996).

In this study, we addressed some of these issues with a reinforcement learning (RL)-based (Sutton & Barto, 2018) computational model of the interaction between visual and self-motion information in a navigation task. Using visual information from a simulated environment and a vector representation of the goal, the reinforcement learning agent learned to navigate in an uncertain environment where either signal is periodically lost. Our modeling results suggest that the nature of the signals and their reliability modulate the interaction of the two signals and how they influence navigation behavior and spatial representations. While we show this here for vector and visual information, the results can be extended to include other potential information sources available to the animal as well.

## Results

We modeled how signals used for making navigation decisions interact with one another and guide navigation using deep RL (Mnih et al., 2015). Our computational model relies on two inputs to navigate to an unmarked goal in the environment — naturalistic visual inputs from the simulation environment in the form of RGB images and a goal vector that is updated at each step. The RL agent receives feedback from the environment in the form of a scalar reward that enables it to learn optimal behavior through trial-and-error. We used a navigation task which we refer to as guidance, where the agent must navigate to an unmarked goal from different start locations. The agent was trained to solve this task under two conditions— one where the agent must learn to use both the visual and vector signal to navigate optimally, and one where either signal or a combination thereof could be used to navigate to the goal.

The crucial element in our model is a feedforward network that processes the two separate input streams, combines them, and makes suggestions for the next action to take (Fig. 1). The final layer of the network consists of action selection neurons, which enable the agent to take translational and rotational actions to navigate in the environment. After it has learned to navigate to the goal, we examine the behavior of the agent and the spatial representations that emerge in the network in order to understand how the agent uses the two input streams to make navigation decisions.

**Figure 1:**
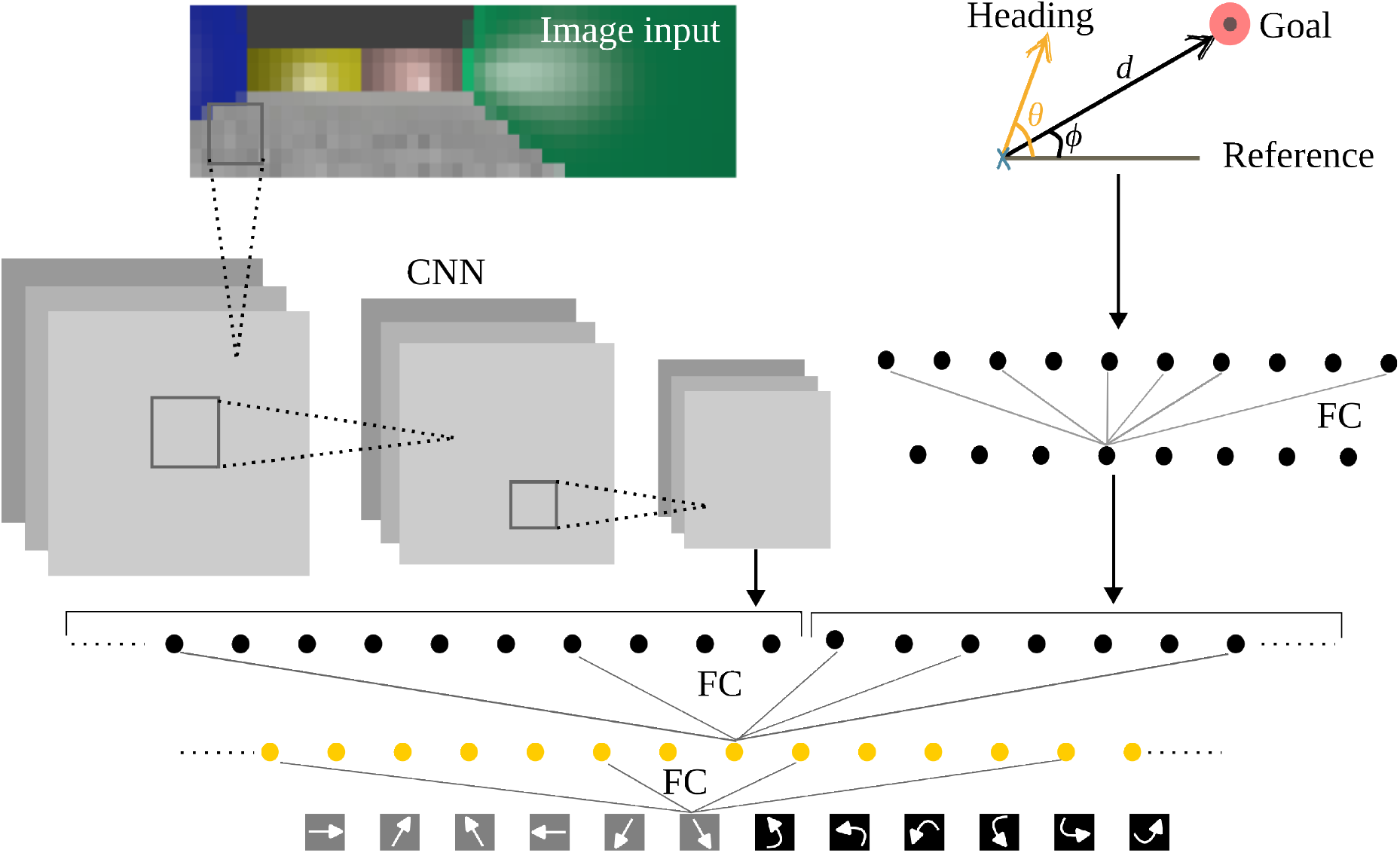
Schematic illustration of model architecture. The network has two separate streams that process visual and vector inputs before integrating them. The visual stream uses a convolutional neural network, and the vector stream uses a fully connected network (FC). For simplicity, only selected connections are shown in all layers. The two streams are then combined and jointly processed. The final layer consists of action units that determine the direction of translation (grey icons) or rotation (black icons) of the agent. Translation actions move the agent by a fixed distance in one of six directions, whereas rotation actions turn the agent to face one of these directions.

### Combining multiple sensory signals aids navigation under uncertain conditions

We first set up a task designed to simulate environmental uncertainty. Humans and animals have to deal with navigating in uncertain environments, where one or more sensory signals may become unreliable or not be available at all. This can be due to several factors, such as poor lighting conditions, occlusion of the field of view by objects in the environment, a lack of external cues that allow for the correction of the path integration signal, or even sensory impairment. Intuitively, being able to use multiple sensory signals adds redundancy and increases the chances of successfully navigating in uncertain environments. For instance, humans rely on a combination of visual, auditory, and proprioceptive cues to maintain balance and navigate through their surroundings (Teaford et al., 2023). The ability to dynamically rely on alternative sensory signals when one becomes unavailable can greatly enhance navigation capabilities in unpredictable environments.

In our task, like the Morris Water Maze (Morris et al., 1982), the agent navigated to an unmarked goal in the environment from various starting points (Fig. 2A). To simulate the temporary loss of a sensory signal, input to either the vector or the visual stream was turned off for 10 trials every 30 trials (Fig. 2B). In this scenario, in order to successfully navigate to the goal on all trials, including those where one signal is lost, the agent had to learn to navigate using each signal independently as well as using them together when both are present. We found that the reinforcement learning agent is able to learn this task efficiently, as demonstrated by the increasing learning curve (Fig. 2C). Additionally, after learning, we assessed the agent’s ability to navigate using either one of the input streams, or both, in a test phase. Test performance shows that the agent is able to navigate to the goal from multiple start locations using either visual or vector streams individually and both together (Fig. 2D). Thus, using and combining multiple sensory inputs lends flexibility and robustness to the agent, allowing it to cope with signal loss and navigate in uncertain environments.

**Figure 2:**
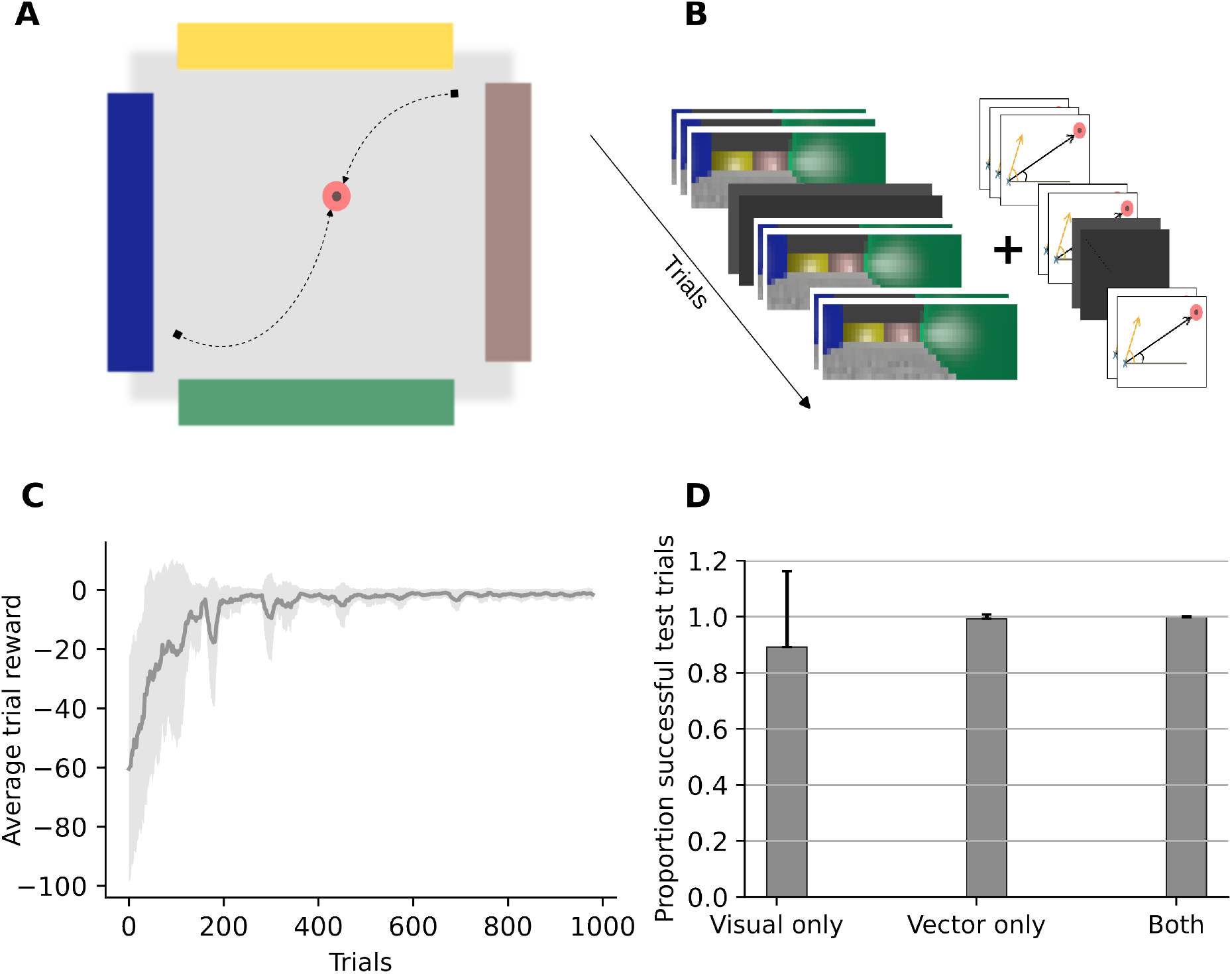
Task structure and model performance. **A**: The task setup for guidance. The agent must navigate to a fixed, unmarked goal from different start locations in the environment. **B**: The task structure. The agent receives vector and visual information through the corresponding network streams at each time step. During training, either stream is temporarily shut off every 30 trials, forcing the agent to learn to use both input streams. **C** : The learning curve shows that the agent is able to learn to navigate well in the uncertain environment task. Light grey area shows the standard deviation. **D** : Performance of the agent in the test phase using visual, vector or both inputs. While test performance with vector or both inputs is nearly indistinguishable, it suffers while using visual input only, indicating a stronger reliance on vector input. Error bars show the standard deviation.

Next, we trained the agent with increasing levels of noise in either the visual or vector stream. We observed that as the noise level increases in either stream, the agent’s direct finding performance gradually worsens, as expected. Interestingly, the performance of the agent is affected more by the addition of noise to the vector stream compared to the visual stream (Fig. 3). We discuss this effect in detail in the following subsections.

**Figure 3:**
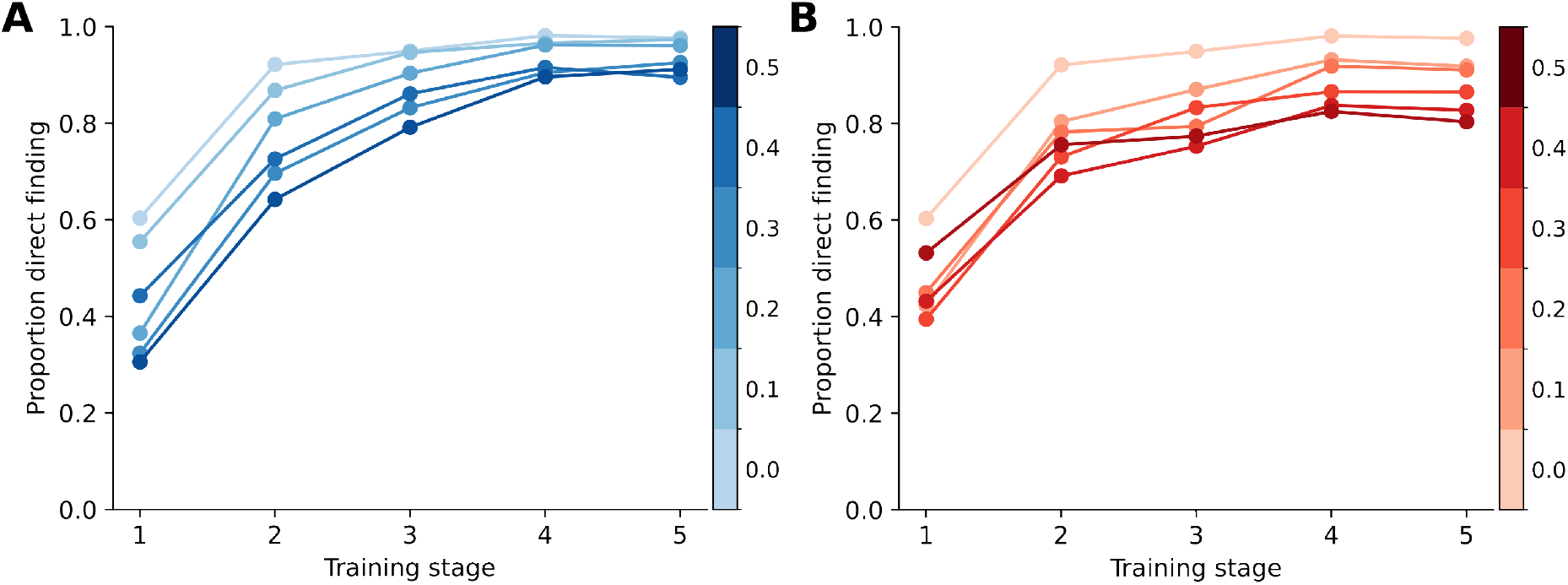
The effect of input noise on spatial learning and navigation performance. In general, increasing the amount of noise in the input streams reduces performance during learning as measured by direct finding. The colorbar represents the variance of the noise added to the input signal added to the **A**: visual stream, or **B**: vector stream. While the effect of input noise in the visual stream is greater at the start of training and becomes smaller as training progresses, the effect of input noise in the vector persists throughout learning.

### Combining multiple sensory signals when a single signal is informative

The results from the simulated uncertain environment demonstrate that being able to use multiple signals to navigate is crucial for robustness in dynamic tasks that are typical for the complexity of real-world navigation. We next study the effect of combining signals in simple tasks when this is not the case, that is, scenarios where a single input stream is perfectly informative and sufficient to navigate successfully. This scenario is typical of most experimental set-ups, where, for instance, vision may be fully sufficient to navigate to the goal. To this end, we again trained the agent to navigate to an unmarked goal in the environment (Fig. 2A), but in contrast to the previous set of simulations, the agent received both vector and visual input in all trials. This way, the agent did not necessarily need to learn to use both streams or integrate them in order to navigate successfully; either stream by itself is sufficient to successfully navigate to the goal.

To examine the factors that drive the integration of the two input streams, we injected noise into either the vector or the visual stream during training while both inputs were available and compared the performance to the cases where the agents received only one input (Fig. 4A, vector only and visual only). The performance of the vector-only agent is more affected by input noise than that of the image-only agent. These differences emerge early on in learning and persist until the final learning phase (Fig. 4A, right panel). Interestingly, although the vector-only agent has the worst direct-finding performance with increasing noise, an agent with noise in the vector stream, but inputs from both streams, navigates far better and, counterintuitively, improves performance with increasing levels of vector noise — performing at the level of a noise-free agent. Such a paradoxical effect is not observed when noise is introduced into the visual stream. While the agent with noise in the visual stream outperforms the image-only agent, the performance of both agents decreases with increasing levels of visual noise, as expected.

**Figure 4:**
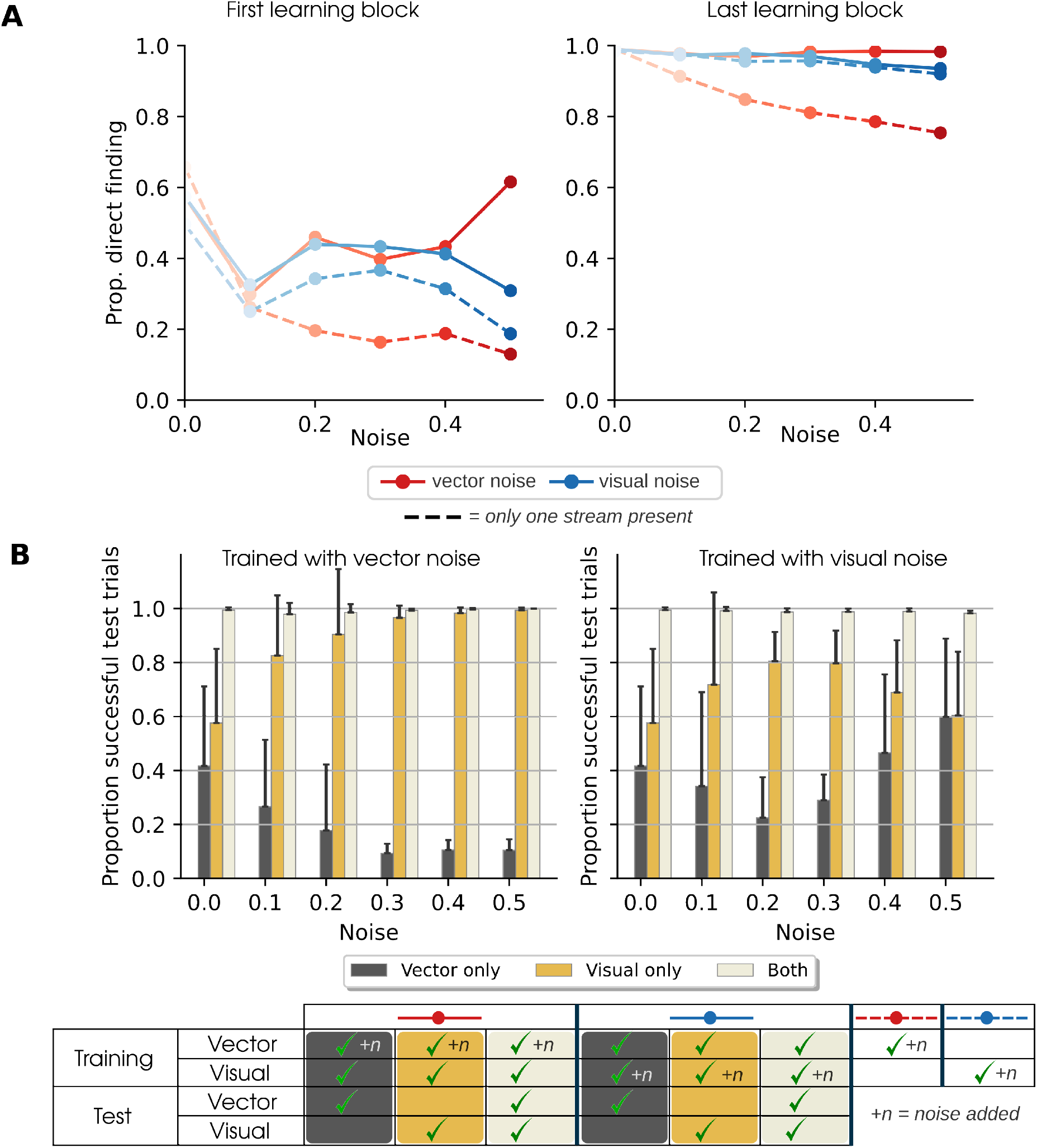
A noisy signal is sometimes ignored rather than integrated. **A:** Direct finding in networks with varying levels of input noise at the beginning (left) and towards the end (right) of learning. Dashed lines correspond to networks trained with both streams active, while the solid lines correspond to networks trained with only one of the two streams receiving input and the other turned off. Generally, increasing noise levels leads to decreasing performance, except in the case when vector noise is added when visual information is also available (dashed red lines). This latter effect appears early on, already becoming apparent in the first learning block. **B:** Test performance of networks trained with varying levels of noise in visual and vector input. During testing, the input sources were entirely reliable, i.e., no noise was added, to assess to what extent the agent learned to use the noisy stream by incorporating it in its navigation decisions, versus having learned to discard it. In the latter case, we would expect there to be almost no difference in test scores between navigating with both streams and navigating without the stream with training noise. With lower levels of noise in the vector stream, the agent still integrates the two sources of information. With increasing vector noise, however, the agent begins to rely solely on the visual stream to navigate, as evidenced by poor test performance when using the vector stream alone, but near perfect test performance when using either both streams or the visual stream alone.

We hypothesized that this difference between agents with noise in the visual and vector streams could be attributed to the degree to which the two signals were integrated to guide navigation, or not. Integrating a noisy signal with a noise-free one may come at the cost of performance, hence pushing the agent towards selecting the noise-free channel. We investigated this potential effect by comparing the test performance of the agents with vector input only, visual input only, and both inputs. The inputs are noise-free in the test phase, allowing us to gauge the degree to which the agent relies on that particular signal to navigate. When the agent is trained with noise in the vector input, it begins to rely entirely on the visual stream to navigate, effectively ignoring the vector input. Thus, as the noise in the vector stream increases, the difference between testing with both inputs (as during training) and testing with only visual input becomes negligible (Fig. 4B, left panel). In contrast, when noise is introduced to the visual stream, the agent still relies on both signals to navigate. When visual input is used to navigate, the test performance is in fact better than when using vector input. Only at greater noise levels does the test performance of the agent with only vector inputs match that of the agent with only visual input (Fig. 4B, right panel). While at first glance the difference between adding noise to one input stream, or the other, may seem surprising, we propose that this has to do with the nature of the signals in question. Images are inherently structural and, even adding noise or occluding a portion of the visual field only weakly interferes with the information in the images that support navigation. On the other hand, adding noise to the vector signal has the potential to substantially deteriorate navigation behavior. If we think of the goal vector pointing toward the goal as being similar to a compass needle pointing north, a small change in the pointing direction could significantly impact navigation success.

In a nutshell, our model highlights the sensitivity of the goal vector to noise and demonstrate that when the vector signal becomes too noisy, the agent begins to ignore this signal and relies solely on the visual signal to navigate.

### Integration of sensory signals explains counter-intuitive experimental findings

Our results so far indicate that using a single stream to navigate when it is sufficient is preferable to attempting to integrate it with another redundant stream, if that stream is noisy. We can use this finding to model two experimental findings in animals with impaired path integration. First, we model results from an experimental study that selectively disrupted the vestibular system of rats, thus presumably interfering with path integration (Stackman & Herbert, 2002). The study found that the number of trials needed to meet a performance criterion (80% correct trials in two consecutive 16-trial blocks) was the same for both sham and lesioned rats when they were trained to find the goal in a Morris Water Maze starting from the same home box position. However, there was a significant difference when the holding box position was changed between trials, with the vestibuar lesion rats surprisingly doing better (Fig. 5A, top).

**Figure 5:**
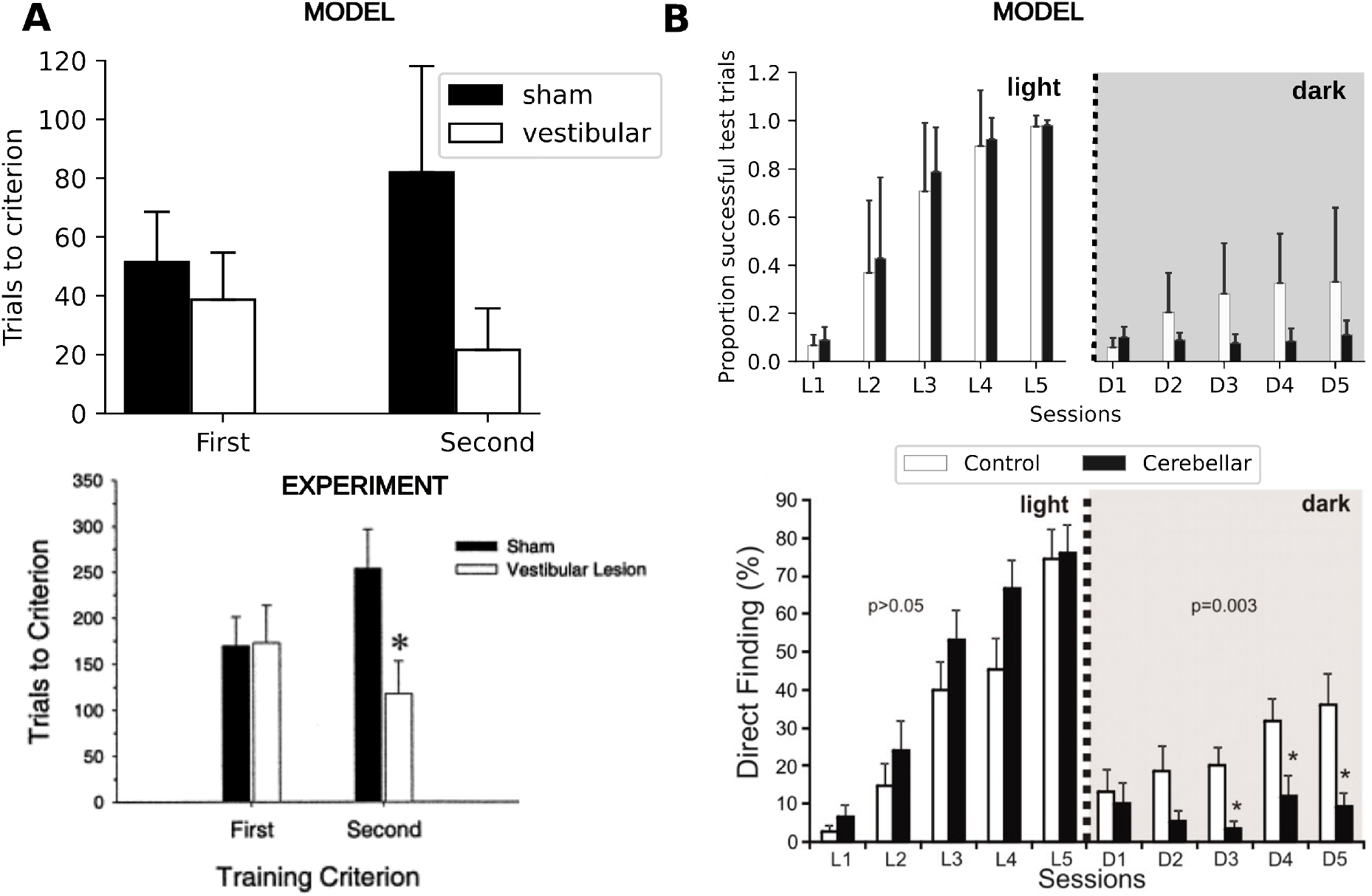
Discarding a noisy vector signal can explain counter-intuitive experimental findings. Modeling results are shown at the top, experimental results at the bottom. **A:** Rats with vestibular lesions found a hidden platform faster than controls did when the holding box was moved between trials compared to control rats (Stackman & Herbert, 2002). Shown are the number of trials to criterion, i.e., lower is better. We used an agent with low vector noise to model the controls and one with the vector input removed for the vestibular lesions (Bottom reproduced with permission from Stackman & Herbert (2002)). **B:** Mice with cerebellar lesions learn to find the goal faster than controls under light conditions, but fail to learn in dark conditions (Rochefort et al., 2011). Shown are fraction of direct finding of the goal, so higher is better. An agent with low vector noise (signals are integrated) is used to model controls, while an agent with high vector noise (vector signal is largely ignored) is used to model the cerebellar mice (Bottom reproduced with permission from Rochefort et al. (2011)).

We used a network with the vector stream removed, i.e., with only the visual stream, to simulate the rats with vestibular lesions, and the regular two-stream network with moderate vector noise to model those with sham lesions. After the agent has learned to navigate to the goal from a single start location, like the rats staring from a single holding box position, we next tasked the agent with finding the goal from four different starting positions, corresponding to four different holding box positions in the experiments. Similar to the experiments, we only see a small difference in performance between the agents when starting from a single start location and a bigger difference in the subsequent phase with four starting positions, with the visual-only network doing better (Fig. 5A, bottom). We propose that the difference in performance between the two agents is small in the first condition where the holding box position remains fixed because i) the task is relatively easy to learn, and ii) a moderately noisy vector signal will still point in roughly the correct direction, which allows the agent to integrate the two signals and correctly navigate from the fixed start position to the goal. In the second condition, where the holding box position is moved between trials, animals with vestibular lesions and the agent with only the visual stream can adapt their behavior quite quickly and build upon the prior knowledge acquired during the first condition to navigate to the goal from different start locations. However, for the sham animals and agents with moderate noise in the vector stream, they must now learn how to integrate the noisy vector information from very different start positions, and the previously learned strategy for integrating the vector signal from a single starting location may actually interfere with learning a general strategy for correctly integrating this signal from any start position, thus affecting performance and slowing down learning of the second navigation condition.

Next, we simulated results from a different experiment in genetically modified cerebellar mice. In the experiments, cerebellar and control mice were tested on their ability to navigate to the goal in a Morris Water Maze under light and dark conditions (Rochefort et al., 2011). We simulate the navigation in light and dark conditions of cerebellar and control mice by again using two different versions of our agent. We liken agents with very high vector noise to animals with an impaired path integration system and use this to model the cerebellar mice. Similar to the previous simulations, we use an agent with moderate vector noise as the control condition since path integration is inherently noisy. As in the experiment, we test both versions of the agent at different stages of learning, either with both visual and vector signals (light condition) or with only the vector signal (dark condition). Our model successfully captures the experimentally observed pattern of direct finding performance at different stages of learning, both in light and dark conditions. When tested under light conditions, both agents improve their direct finding performance over time, similar to mice in the experiment. In addition, our model also explains the small but consistent advantage of the cerebellar mice in the light condition. Our model accounts for this finding based on the fact that agents with high noise in the vector stream learn to ignore the noisy signal quite early on in learning in navigation tasks that have reliable visual input, while models with moderate noise still integrate both signals and take longer to learn this integration. This slight advantage, however, comes at the cost of robustness, as evidenced by the poor performance of the model with high vector noise and cerebellar mice when tested in the dark.

### Spatial representations are differently affected by the removal of sensory input

We next set out to understand the types of spatial representations that emerge in the agents’ networks to support navigation and how the representations are modulated by visual and vector inputs. Place cell activity is minimally disrupted in the dark in rats (Quirk et al., 1990), and in congenitally blind rats (Save et al., 1998). On the other hand, place cell activity is more dramatically disrupted by removing vector input (Stackman et al., 2002; Rochefort et al., 2011).

After training the network on guidance in an uncertain environment, we first computed spatial activity maps for all the neurons in the layer prior to the action selection layer and classified them (see Methods). A number of place-like units emerged in the network (Fig. 6), consistent with previous work showing that an agent learning to navigate to an unmarked goal using visual landmarks tends to develop place-cell-like representations to support navigation (Vijayabaskaran & Cheng, 2022).

**Figure 6:**
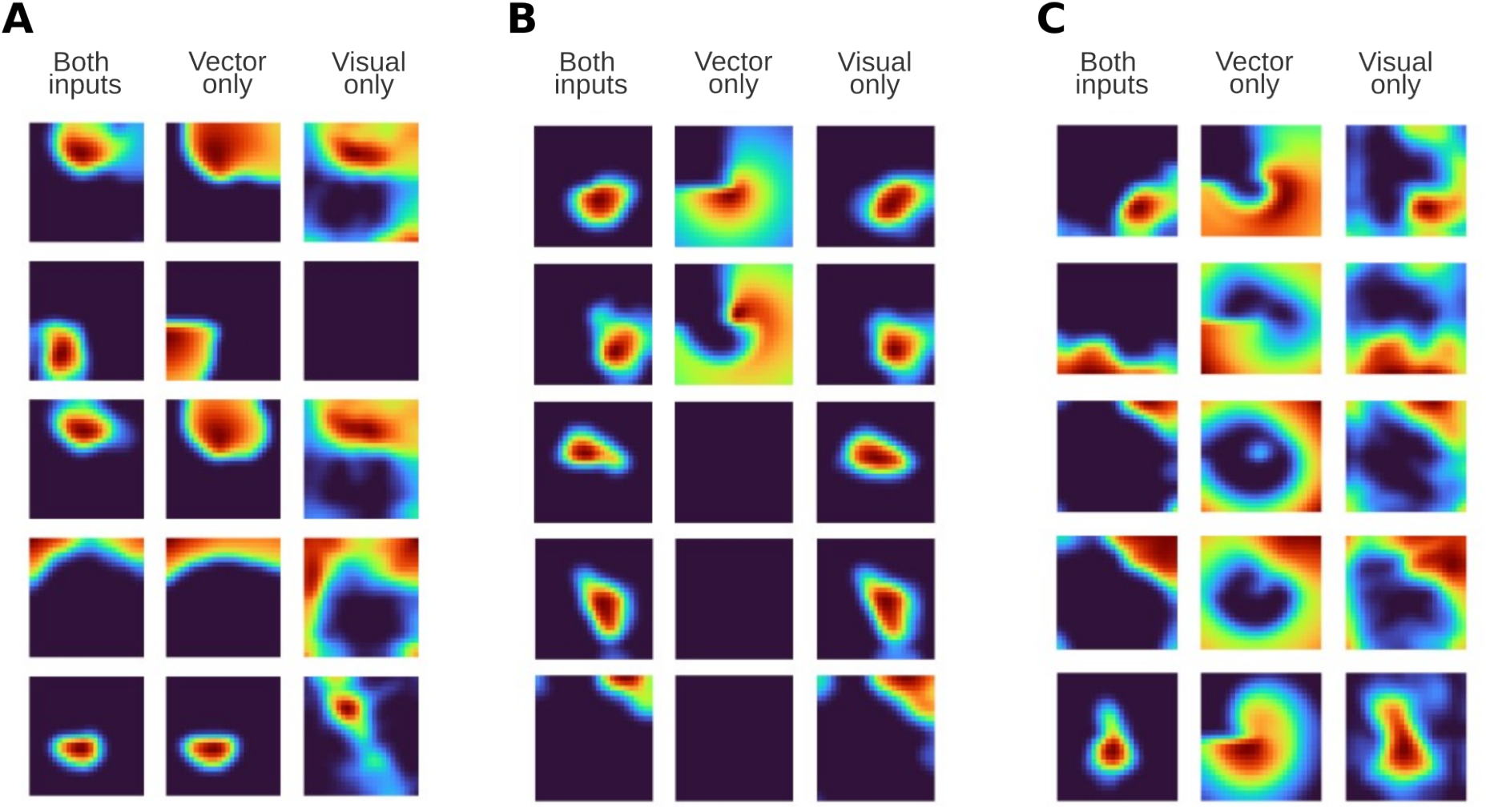
Place cell representations have a complex dependence on visual and vector inputs. Place cell representations emerge in the agent as a result of learning to solve the task. When either visual or vector input to the network is removed, remapping occurs to different degrees (see also Fig. 7). Example fields that **A**: remain place-like with vector input only, **B**: remain place-like with visual input only, and **C**: need both inputs to remain place-like.

We next studied the effect of removing either visual or vector input on the place cell representations by recomputing the spatial activity maps with either one of the inputs removed. We found that removing either input results in some disruption of place-like units in the network; some of these units require both inputs to maintain their firing field, while others are able to maintain their firing field in the presence of one input alone (Fig. 6). Removing the vector input results in a larger drop in the number of place cells than removing visual inputs (Fig. 7A), consistent with experimental reports that place field firing is largely maintained in the dark (Quirk et al., 1990), but disrupted when the vestibular system is disturbed (Stackman et al., 2002).

**Figure 7:**
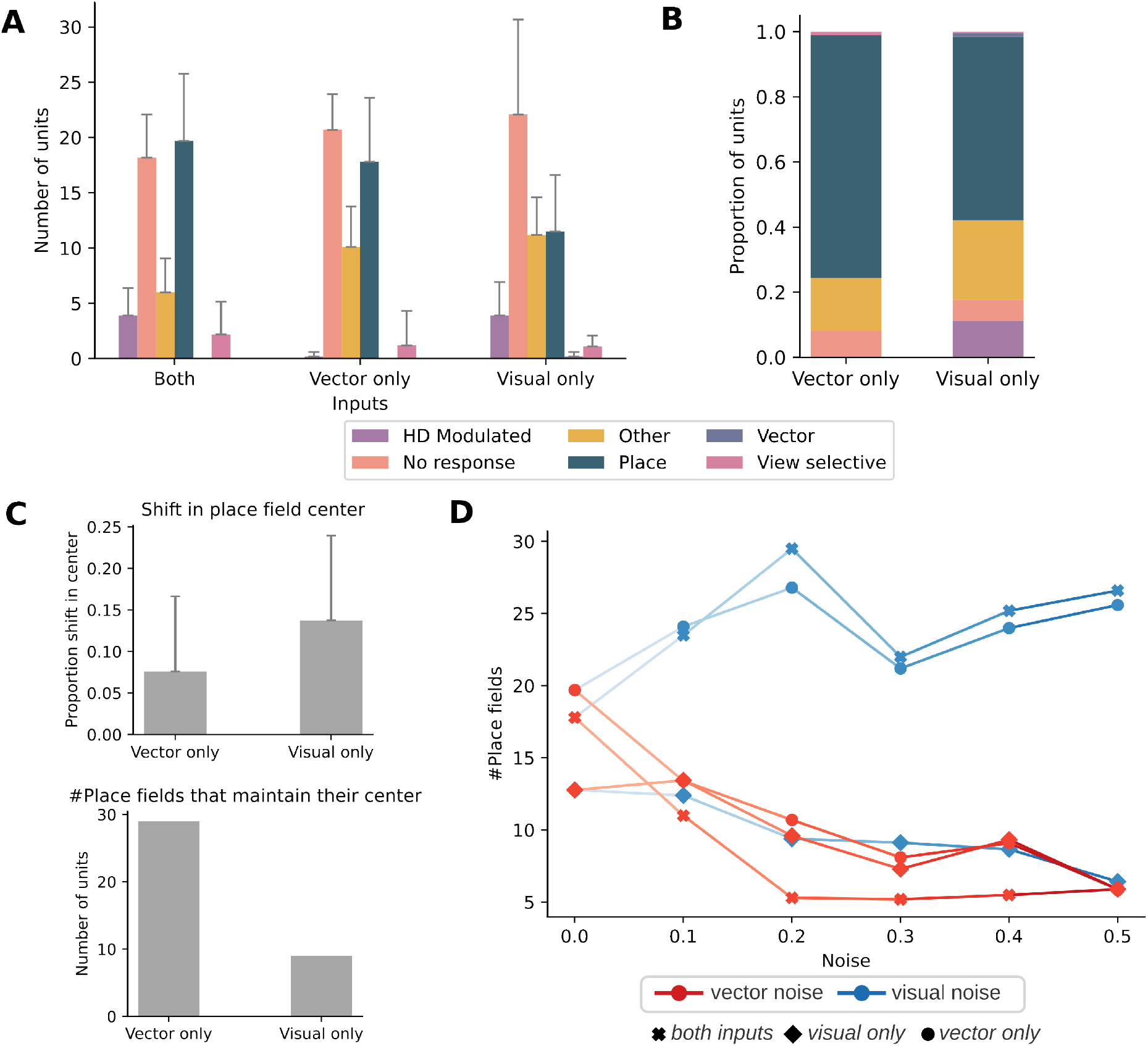
Spatial representations remap when inputs are removed. Removing either visual or vector input disrupts place cell representations. On average, this effect is more pronounced when vector inputs are removed (visual only). **A**: Distribution of response types in a network with both inputs, and with only one of the inputs. There are fewer place-like units when the network receives only visual inputs, i.e, after the removal of vector input. **B**: Redistribution of place cells after removal of an input. The majority of place cells continue to remain place-like. The place cells that change to other response types typically either completely lose their response (silent), indicating that their firing was primarily driven by one input type, or the firing field loses its locality (“other”). **C**: The effect of training noise on the number of place-like units. The influence of noise on the number of place fields is determined by the signal to which the noise is added during training. When there is noise in the visual stream, the total number of place-like units is not affected much by the addition of noise, but they become increasingly driven by the vector stream. Hence, removing the vector stream leads to a drop in the number of place-like units. On the other other hand, increasing the noise in the vector stream reduces the number of place-like units irrespective of which input is removed. These results correspond to the higher effect that noise in the vector stream has on direct finding performance. (Fig. 4) **D**: Remapping of place-like units. On average, of the place-like units that remain place-like with only one input, more units maintain their firing field center (top) with lower shifts in firing field center (bottom) when there vector input than with visual input.

We also observed that the number of units classified as “other,” which means that the units do not have a localized firing field, as well as the number of silent cells, increase with the removal of either input, suggesting that place-like units either become silent or lose their local firing properties with the removal of one input. To study the fate of place-like representations more systematically, we examined what happens to place-like units after the removal of either input (Fig. 7B). In both cases, more than half of the units remain place-like even after the removal of one input. As suggested by the previous result, the remainder of the units mostly either lose their local firing properties or become silent. In addition, a significant proportion of cells become head-direction modulated upon the removal of the vector input. That is, these units continue to have local firing fields, but the location of the firing field is now modulated by the heading direction.

Since the majority of the place-like units remain place-like even after the removal of one input, we next ask if those units maintain the location of their firing field or if they remap to new locations, and if so, to what extent. We first quantify the proportion of place cells that maintain their firing field after the removal of input. Most place cells continue to maintain the same firing field location with vector input only (Fig. 7D, top), much like place cells in rats navigating in the dark. Fewer cells show the same property with visual input only. Next, for the cells that do not maintain their firing field, we also compute the average shift in the location of the firing field (Fig. 7D, bottom). This also reveals that for those cells that do show a shift in the center of their firing field, the removal of visual input tends to show a lower average shift in place field center as compared to the removal of vector input. Taken together, these results suggest that in our model place field firing is modulated by both visual and vector inputs and that, overall, the removal of vector input results in a greater disruption of firing field locality and more remapping of place-like units.

Finally, we also looked at the effect of input noise on the number of place-like units in the network (Fig. 7C). First, we find that adding noise to the vector input drastically reduces the number of place cells in the network. With a noisy vector input, removal of the vector input has a negligible effect on the number of place cells in the network, which is consistent with the results in Fig. 4 that show that with a noisy vector input, the agent begins to increasingly rely on the visual input to guide its actions. In contrast, with noise in the visual stream, removing the vector input has a large negative impact on the number of place cells in the network. However, removing the visual input has a negligible effect on the number of place cells; in fact, as noise is injected into the visual input, the number of place cells even increases slightly when the noisy input is removed. This suggests that a small number of units that are simultaneously driven by vector and visual input, but whose localized firing is disrupted by the noisy visual stream, recover their locality when this input is removed.

Taken together, the analysis of spatial representations in the network paints a picture similar to that of the behavioral analysis: place-like representations are driven by both visual and vector input, with vector inputs playing a more vital role in driving them.

### Realistic implementation of the vector stream

In our simulations so far, the model had access to an already computed goal vector, which we assumed is the result of a separate computational process such as path integration. While path integration using self-motion is the best candidate for updating the goal vector, it could in practice be acquired from another process, for example, from path integration using optic flow, visual estimation, or an existing metric map of the environment. Animals could also use one of the above methods to occasionally re-calibrate and correct errors from path integration, or use some combination of the above. We next explored two simple methods using a goal vector that is learned from visual information instead of hard-coded. In the first method, we used a supervised learning network module to estimate the goal vector from visual inputs (Fig. 8A). There is evidence that humans and animals can estimate the goal vector visually (H.-J. Sun et al., 2004; Lappe et al., 2007). In the second method, we use the same network structure as the supervised learning task, but this time the network is attached to the network shown in Fig. 1, and this entire, larger network is trained end-to-end with reinforcement learning.

**Figure 8:**
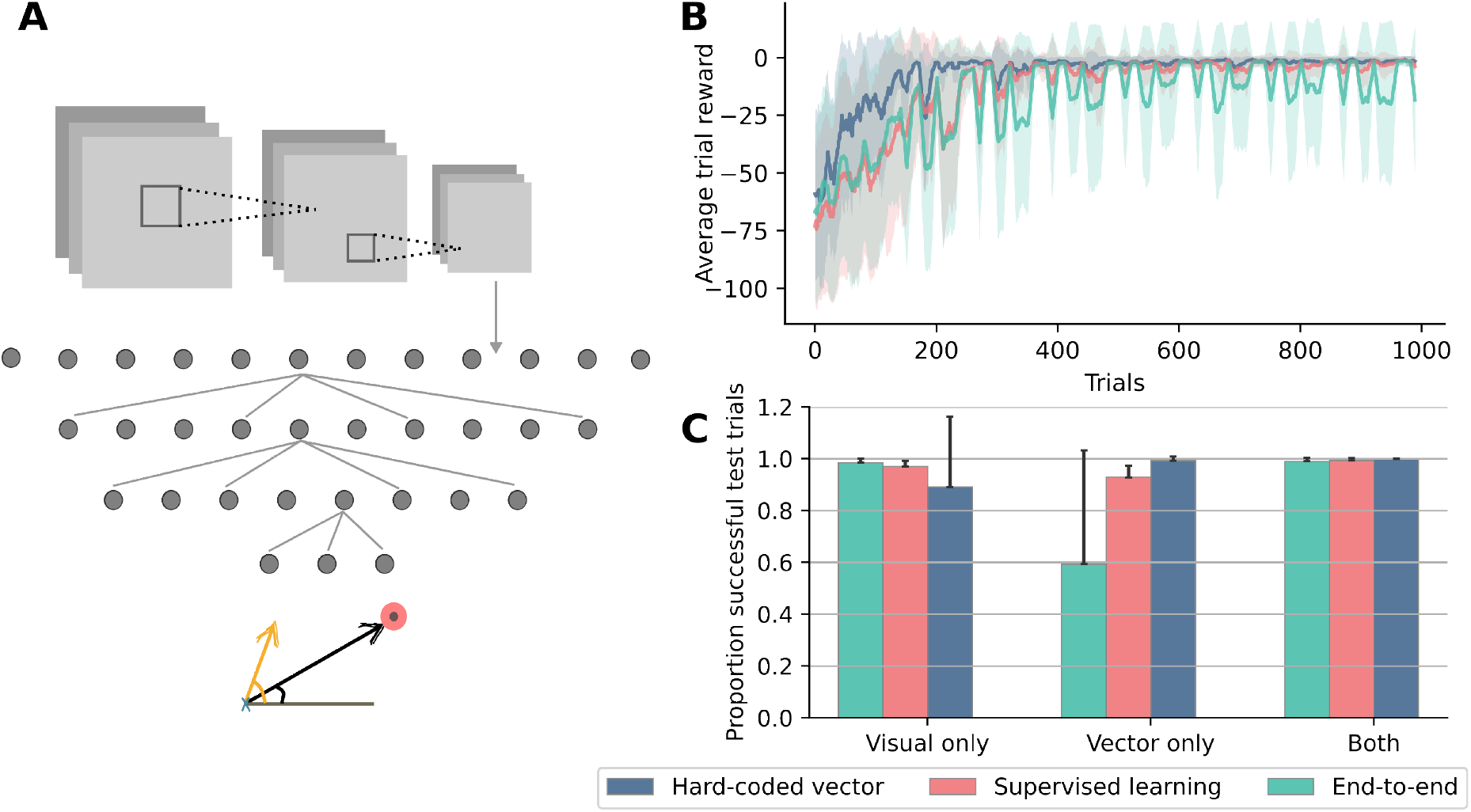
The effect of the vector encoding scheme on learning and performance. **A**: The network used to learn a vector representation. The network is either trained end-to-end as part of the vector stream depicted in Fig. 1A using reinforcement learning, or it is pre-trained to predict the goal vector using supervised learning. **B**: Learning curves using the three vector encoding schemes. The network successfully learns the task using all three schemes, however, learning is slower and noisier for the end-to-end scheme and is fastest while using a hard-coded vector. **C**: Test performance of the three vector encoding schemes measured with both inputs, and using either one of the inputs only. While the performance for all three schemes is comparable when both inputs are available, the reliability of the vector encoding scheme determines how well the network performs when only vector information is available. The better the vector encoding, the worse the network performs when only the visual stream is active.

We trained these two variations of the model in the uncertain environment task (Fig. 2B) and compare it to the standard model that uses a hard-coded vector. Examining the learning curves shows that both variants can successfully learn the task, with the model trained end-to-end via reinforcement learning displaying slower and noisier learning than the others (Fig. 8B). We also tested the two variants, along with the hard-coded vector model, for comparison with each input separately. Consistent with the results from studying the noisy models, we find that the noisier the vector encoding, the better the network performs using visual input alone (Fig. 8C).

Both of the alternate methods discussed in this section use visual estimation to estimate the goal vector. It is of course possible to instead use some type of recurrent network trained via supervised learning to do path integration (Cueva & Wei, 2018; Banino et al., 2018) and use that to predict the goal vector at every step, but this is out of the scope of the current study.

## Discussion

We presented a normative model of the interaction of goal vector and visual information for navigation based on deep reinforcement learning. We found that in general, the agent is more sensitive to noise in the vector stream, and that whether the signals are integrated or one is chosen over the other is primarily modulated by the amount of noise in the signals. Integrating signals results in more robust behavior, which is useful while navigating under uncertainty, but can sometimes prove to be disadvantageous in other situations when the discrepancies between the two streams are too large. In the latter case, it might be better to ignore one input altogether.

### Factors affecting the combination of information sources

Our simulations reveal that navigating using multiple sources is a complex and multifaceted problem, with environmental uncertainty, the amount of noise in each signal, and the nature of the signal itself playing a role. Here, we discuss in more detail each of these factors and their impact on navigation behavior. Environmental uncertainty was incorporated into our simulations by intermittently removing either the visual or goal vector input during navigation, and we find that the agent is able to cope well with these dynamic changes. In experiments, humans and animals have also shown the ability to adapt well to the removal of sensory inputs. Gerbils navigating to food locations using an array of landmarks still reached their goal when the light was cut off mid-trajectory (Collett et al., 1986). Similarly, rats trained to solve the Morris Water Maze were able to navigate accurately in the dark after starting their trajectory in the light (Save, 1997), and blindfolded humans can navigate to a target that they had seen prior to being blindfolded (Mittelstaedt & Mittelstaedt, 2001). While removing visual input in experiments is fairly simple to achieve by blindfolding or turning off lights, stopping path integration and the estimation of a goal vector can be quite challenging. One alternative is to temporarily lesion structures necessary for path integration. For instance, Stackman et al. (2002) find that rats with lesions to their vestibular apparatus show the ability to navigate accurately to a goal when visual cues are present, but fail to do so in the absence of these cues, while those with sham lesions have no difficulty doing so. This technique, however, impairs path integration and, thus, the goal vector update for the entire duration of navigation, not mid-navigation like our simulations. Even so, it is likely a reasonable modeling assumption that the goal vector will not always be available during navigation. This is due to the fact that accumulating noise can easily degrade the path integration signal, which necessitates periodic resets by environmental boundaries or recognizable landmarks (Mizumori & Williams, 1993; Goodridge et al., 1998). Thus, environmental uncertainty is an important factor that influences how signals interact to inform navigation decisions.

Next, the amount of noise in each information source also affects how signals are combined. In the model, depending on how much noise is present, a signal is either integrated with the other, more reliable signal or ignored. This observation is closely related to those from studies that aim to identify whether signals cooperate or compete in order to determine movement direction. This is typically accomplished by bringing visual and vector (path-integration) signals into conflict with one another. Multiple studies have shown that signals are integrated for low and moderate mismatches and compete for higher mismatches (Etienne et al., 1996; Etienne & Jeffery, 2004; X. Chen et al., 2017; Zhao & Warren, 2015; Foo et al., 2005), which is consistent with our finding that for lower noise levels, the two signals are integrated (cooperation), but at higher noise levels one signal is selected over the other (competition).

Finally, our model demonstrates that in cases of competition, it is not only the mismatch that determines which signal is selected but also the nature of the signal. Since visual information is more structured and hence robust to noise, when there is a conflict, it is preferentially selected over vector information. Similar observations have also been made in experiments (Etienne et al., 1998). This suggests that competition between signals is resolved by a hierarchical organization of preferences. Indeed, it has been shown that in rats, there is a hierarchical organization of visual, odor, and vector information (Maaswinkel & Whishaw, 1999). Taken together, our results provide an account of the integration of signals that is neither purely cooperation nor competition but rather a hybrid of the two mediated by the factors discussed above.

### What drives spatial representations?

While the primary focus of this study was on behavior, our findings can also add to the current understanding of what drives spatial representations in the brain. Place cells and other spatially selective cells do not exist in a vacuum, as previous research has shown; rather, the nature of the task at hand and the navigation strategy used both interact to shape spatial representations (Vijayabaskaran & Cheng, 2022). Our model further extends this idea by demonstrating that sensory inputs and their primacy and reliability can also influence spatial representations. This is in line with several studies showing that path integration signals (Gothard et al., 1996; Redish et al., 2000; Bjerknes et al., 2018; G. Chen et al., 2013), taste (Herzog et al., 2019), odor (Fischler-Ruiz et al., 2021; Zhang & Manahan-Vaughan, 2015; Save et al., 2000; Aikath et al., 2014; Radvansky & Dombeck, 2018), auditory (Moita et al., 2003), haptic (Gener et al., 2013), and visual cues (Muller & Kubie, 1987; G. Chen et al., 2013; O’Keefe & Burgess, 1996) all affect hippocampal place cell maps. Specifically, with regard to vision and self-motion, modeling and experimental work have shown that vision alone is sufficient to drive place representations in a smaller proportion of cells (G. Chen et al., 2013; Vijayabaskaran & Cheng, 2022). Our current study suggests that the dependence of place-like spatial representations on each signal is heterogeneous, with a greater reliance on vector input to maintain the firing field. This is qualitatively in agreement with G. Chen et al. (2013)’s study in virtual reality, which found that overall, vision alone was sufficient for 25% of CA1 place cells, with the remaining 75% requiring input from self-motion, although to varying extents.

Different species will rely on various sensory inputs to varying degrees, so we might anticipate variations in the kinds of spatial representations and behavioral performance based on the primacy and reliability of various sensory signals. Indeed, in species with a highly developed visual system, such as primates, hippocampal cells respond less often to a particular location and more commonly to a particular view (Rolls, 2023).

Finally, further complicating the issue, many hippocampal cells exhibit mixed selectivity for a variety of additional, non-spatial, task-related variables, including lap number in a multi-lap run (C. Sun et al., 2020), route taken (Wood et al., 2000; Grieves et al., 2016), position of another animal (Smith & Mizumori, 2006; Sarel et al., 2022), accumulated evidence for a choice (Nieh et al., 2021), and frequency of sound (Aronov et al., 2017). Thus, teasing apart exactly what factors drive spatial representations and to what extent will require further modeling and experimental work.

### Limitations and relationship to other models of multisensory integration

How to best combine and use information from multiple sensors in order to guide decisions is a question that has also been widely explored in engineering, notably in robotics. This problem, known as sensor fusion, is still an area of active research and development, with much effort being focused on improving the accuracy and reliability of navigation in mobile robots (Soloviev & Miller, 2012). Commonly used methods include Kalman filters and Bayesian approaches, among others, and are often inspired by sensory integration in nature (Murphy, Jan./1996). Bayesian frameworks have also been used to model multisensory integration in humans and animals (Knill & Richards, 1996) and, in general, provide an account in which signals are combined by an weighted summation process, based on the system’s belief in how reliable the components are. This idea has also been extended to include situations where relying on a single signal rather than combining multiple signals may be advantageous (Cheng et al., 2007). Such an extended Bayesian framework shares certain similarities with our model, namely, the emphasis on the reliability of signals and the potential advantage of relying on a single signal. Also related to Bayesian models is the hybrid model of Harootonian et al. (2022), which has two components: a path integration system and a Kalman filter that independently estimate heading direction from idiothetic and allothetic information. Fitting the model to human behavior revealed that a hybrid strategy consistent with combining inputs when the mismatch is small and selecting one when the mismatch is large best described the behavior of all the participants. While our model shares some similarities with the methods discussed above, it does not require the assumption of *a priori* knowledge about which signals are reliable, and the hierarchy of vision over vector information emerges during the learning process. It also goes beyond Bayesian models in that it operates in a closed loop, allowing us to study the interaction between sensory inputs, behavior, and internal representations in the network.

This operation in a closed loop is possible in our model due to the use of RL, which allows the agent to continuously learn and adapt its behavior based on feedback from the environment. However, this approach has its limitations; for example, learning occurs on very different time scales in RL models and in animals. The former often requires several thousands of trials to learn navigation tasks that would take mammals only a few training blocks. Moreover, the types of networks used for learning in deep RL algorithms are not entirely biologically plausible. Regardless of these limitations, we believe that RL can be a useful modeling tool, especially for studying spatial navigation, because of some shared commonalities. For example, both the navigation problem and RL are goal-driven. In addition, they must also deal with balancing exploration and repeating known routes, and must be able to deal with uncertainty. So, while specific features of RL may not match animal learning, we believe that it is a useful tool to understand the computational problem at hand and allows the exploration of different elements that contribute to navigation, such as the perception of sensory inputs, memory and replay, and decision-making. Moreover, RL models can be tuned and manipulated to explore various scenarios and test specific hypotheses. Making use of a neural network also enables the use of natural sensory representations (such as visual input) and the analysis of spatial representations that emerge in the network to support behavior. Together, these can provide valuable insights into underlying mechanisms and help uncover general principles governing different aspects of navigation.

In summary, a trained RL agent in our model can be seen as a normative model of how an agent ought to combine different sources of information to achieve maximum performance in spatial navigation. Our simulation setup allows us to study the optimal strategies for information fusion in realistic settings while at the same time revealing emergent spatial representations that enable it to do so. The modeling results are consistent with experimental results and frequently defy simple intuitions about the effects of removing a sensory input on neural spatial representations, and spatial learning and navigation.

## Methods and materials

### Task structure

Unless otherwise stated, all simulations used a simple navigation paradigm, called guidance, which is similar to a dry version of the Morris Water Maze. The task involved navigating to a fixed, unmarked goal location in the environment from different starting locations. At the start of each trial, a start location was assigned at random. To aid in localization, the walls of the enclosure were marked with distinct colors, which acted as distal landmarks. We used CoBeL-RL (Closed-loop simulator of complex behavior and learning based on reinforcement learning and deep neural networks) (Diekmann et al., 2023) to generate the simulation environments and train the agents. The agent received visual input and goal vector information from the simulation environment at each time step. We defined two task schemes to evaluate the agent. In the first scheme, the visual or vector stream was randomly turned off or on every 30 trials, requiring the agent to learn to navigate using both streams simultaneously as well as separately. In the second scheme, both streams were available continuously, such that the agent could freely choose to select a single stream or combine them in order to navigate.

Training lasted for 4000 trials in total and each trial lasted 100 time steps, unless the trial was terminated earlier by correctly locating the goal. The agent received a +1 reward for reaching the goal. The agent received a -1 penalty (negative reward) for each step taken in order to motivate it to find shorter routes. If the agent selected an action that would have resulted in walking into a wall, it remained in place and received a -1 penalty.

### Network model

The network model that was trained to solve the task is shown in Fig. 1. The network had two input layers that receive image and vector information, respectively. The two inputs were processed in separate streams before being combined. The visual stream processes visual information from the environment using a convolutional neural network (CNN) (Lecun et al., Nov./1998). The CNN consisted of three layers with 32, 64, and 64 filters each, with kernel sizes of (5,5), (4,4), and (3,3), respectively. The vector stream received the goal vector through its input layer, which was connected to two fully connected feedforward network layers, which have 50 and 25 units, respectively. The two streams were then combined by a simple concatenation and further connected to a fully connected (FC) layer with 50 units and a dropout of 0.3, before an action selection layer with 12 units, which selected rotational and translational actions. A ReLU activation function (Nair & Hinton, 2010) was used throughout the network, except for the output layer, which used a linear activation function.

The actions available to the agent for interacting with the environment corresponded to the units of the action selection layer. Six rotational and six translational actions were available to the agent, which were aligned with the main axes of a hexagonal grid. Translational actions caused the agent to move to a neighboring node on the grid, while rotations caused the agent to change its heading direction to face a neighboring node.

### Visual and vector inputs

The visual stream received naturalistic RGB images that correspond to the agent’s current view of the environment. Each of the 48 × 12-pixel images corresponded to a 240-degree field of view in the simulated environment. The vector stream received information about the goal vector. The input to this stream is (*d, ϕ, θ*), where *d* is the distance to the goal, *ϕ* is the angle between the goal vector and a reference vector, and *θ* is the allocentric heading direction of the agent with the same reference vector.

The goal vector can also be learned. We demonstrated this using two alternative versions of the model (Fig. 8). In the first version, the goal vector was learned from visual impressions of the environment by a neural network that was pre-trained using supervised learning. The network had the same convolutional neural network as the main model, followed by fully connected layers with 50, 25, and 12 units, respectively, and an output layer with 3 units corresponding to *d, ϕ*, and *θ*, which were used as input to the vector stream of the network depicted in Fig. 1. The second version used an identical network, except that the model was trained end-to-end using reinforcement learning (RL). The details of the RL algorithm are described in the following subsection.

### Reinforcement learning

In RL, the model, referred to as an agent, interacts with its environment and receives feedback in the form of rewards or penalties based on its actions during training, and uses this to optimize the expected cumulative reward. This allows it to learn optimal behavior through trial-and-error — the agent learns from its own experiences and adapts its behavior accordingly. The RL algorithm used in our model is based on Q-learning (Watkins & Dayan, 1992). In Q-learning, the agent learns to estimate Q-values, i.e., the value of each action *a*_*t*_ in a given state *s*_*t*_. These estimates are updated after each action using the following rule:

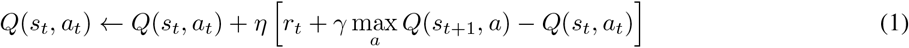

We used the Deep Q-Network (DQN) algorithm (Mnih et al., 2015) to train the previously described network to navigate using visual and vector inputs. We used a learning rate *η* = 0.001 and a discount factor of *γ* = 0.99. The agent drew from an experience replay (Lin, 1992) buffer of 3000 past experiences to learn from. In general, reinforcement learning agents must deal with what is known as the exploration-exploitation trade-off, which refers to the fact that agents must explore new actions to discover better strategies while also exploiting the actions that have yielded higher rewards in the past. We used a commonly employed policy, *ε*-greedy, with *ε* = 0.3 during training. In the test phase, the agent used a greedy policy, i.e., always picked the learned optimal action at every step.

### Behavioral analysis

We evaluated the behavior of the agent both during training, as well as after, in a test phase. A standard way to evaluate agent training performance is to examine the learning curves. In addition to this, in order to get a more fine-grained view of the agents’ performance, we defined a learning phase of 1000 trials in which learning converged in all models. We divided these into five learning blocks of 200 trials each, and calculated the fraction of direct finding trials in each learning block. The performance of the agents was evaluated after training, in a test phase. The agent was tested in three different modes to assess which signals the agent had learned to use to solve the task. In the first mode, the agents received both the visual and vector signals, while in the other two modes, they received only one of the two signals for the entirety of the test phase. During testing, no noise was added to the visual and vector signals. The fraction of test trials in which the agent successfully navigated to the goal was used to measure performance.

### Simulation of cerebellar mice in light and dark conditions

We simulated the experiment described in Rochefort et al. (2011) using the Morris Water Maze task described above. If mice with cerebellar lesions have impaired path integration, we assume that this leads to a highly imprecise calculation of the goal vector. We simulated this by using agents with a highly noisy (variance = 0.8) vector signal. For the control mice, we used agents with a lower level of noise (variance = 0.1) in the vector stream. We do not use networks with zero noise since path integration is subject to accumulating errors and is thus inherently subject to some noise. We trained the networks for 1000 trials each and partitioned the learning phase into 5 learning blocks of 200 trials each. At the end of each learning block, the agents were tested under light and dark conditions. For the light condition, both the vector and visual stream were active, and for the dark condition only the vector stream was active. Each test phase lasted 100 trials, and the proportion of successful trials was computed.

### Simulation of rats with vestibular lesions

We hypothesized that lesions of the vestibular system lead to impaired path integration and simulated this using agents with the vector stream removed and compared their performance to control agents with low noise (variance= 0.1) in the vector stream. Two variations of the task were used, corresponding to the first and second criteria defined in the experiment by Stackman & Herbert (2002). In the first variation, the agent always started from the same location, corresponding to a fixed holding box position in the experiment. In the second variation, four starting locations were used, again corresponding to the holding box positions. To enable comparison to the experimental results, we used an analogous performance measure. The criterion was defined as successful direct finding in three consecutive blocks of 16 trials, and the number of trials to meet the criterion during training was used to quantify the speed of learning.

### Analysis of spatial representations

We calculated spatial activity maps for each unit in the layer before the action selection layer by recording unit activation at each grid point on a 25 × 25 grid on the environment. The maps are smoothed using a Gaussian filter with a standard deviation of 2. This process is repeated for each of the six head directions, and the units are then classified as place cells, egocentric vector cells, head-direction modulated cells, and view-selective cells as outlined in Vijayabaskaran & Cheng (2022).

## Acknowledgments

This work was supported by the Deutsche Forschungsgemeinschaft (DFG, German Research Foundation) – project number 316803389 – through SFB 1280, project A14.

